# Transcription factor LHX2 suppresses astrocyte proliferation in the postnatal mammalian cerebral cortex

**DOI:** 10.1101/2025.04.10.648067

**Authors:** Archana Iyer, Reanne Fronteiro, Prachi Bhatia, Sanjna Kumari, Amrita Singh, Jiafeng Zhou, Riccardo Bocchi, Rishikesh Narayanan, Shubha Tole

## Abstract

In the developing cerebral cortex astrocytes arise from progenitors in the ventricular and subventricular zones (V-SVZ), and also from local proliferation within the parenchyma. In the mouse neocortex, astrocytes that occupy upper versus deep layers (UL/DL) are known to be distinct populations in terms of molecular and morphological features. Transcription factor LHX2 is expressed both in V-SVZ gliogenic progenitors and in differentiated astrocytes throughout development and into adulthood. Here we show that loss of *Lhx2* at birth results in an increased astrocyte proliferation in UL but not the DL of the cortex in the first postnatal week. Consistent with this, transcriptomic signatures of UL astrocytes increase. By 3 months, *Lhx2* mutant astrocytes display upregulation of GFAP, and transcriptomic signatures associated with “reactive” astrocytes, in the absence of injury. These results demonstrate a novel role for *Lhx2* in regulating proliferation and molecular features of cortical astrocytes.

**Summary Statement:** Loss of *Lhx2* causes increased astrocyte proliferation in the upper but not deep layers of the mammalian cortex, and upregulation of reactive gliosis-like signatures in the absence of injury.

## Introduction

Astrocytes, originally considered to be glue-like cementing material in the brain (Somjen, 1988), are now known to play critical roles in establishing and modulating functional circuitry. They mediate neuronal synapse formation and function (N. J. Allen & Eroglu, 2017; Stogsdill et al., 2017; Tani et al., 2014). Astrocytes also modulate blood vessel growth and branching (Ma, Kwon, & Huang, 2012; Tabata et al., 2022), regulate blood flow in a neuronal activity dependent manner (MacVicar & Newman, 2015; Otsu et al., 2015), and modulate nutrient uptake via their processes that wrap around blood vessels (Barres, 2008). In injury response in the central nervous system, astrocytes become reactive and hypertrophic (Escartin et al., 2021)

Despite these diverse functions, astrocytes were initially thought to be a largely homogenous population defined by the presence of the Glial Fibrillary Acidic Protein (GFAP) (Brenner, Kisseberth, Su, Besnard, & Messing, 1994). Morphological distinctions hinted at some non-uniformity, since fibrous astrocytes, found in the cortical white matter, are GFAP+ whereas protoplasmic astrocytes found in the cortical grey matter, are largely GFAP- (Sofroniew & Vinters, 2010). Recent advances using high throughput transcriptomics have revealed a remarkable molecular diversity in astrocytes residing in different parts of the central nervous system, suggesting there may be distinct sub-groups (John Lin et al., 2017; Morel et al., 2017). In particular, astrocytic sub-populations vary in terms of their proliferative nature and their ability to facilitate neuronal synaptogenesis (John Lin et al., 2017; Morel et al., 2017). Within the cortex, astrocytes display cortical layer-specific morphological and transcriptional differences (Bayraktar et al., 2020; Lanjakornsiripan et al., 2018). Here, we show that transcription factor LHX2, an established regulator of neurogenesis and neuronal cell fate, also controls the proliferation of a subset of astrocytes residing in the upper layers of the cortex.

One common feature seen across astrocytic sub-groups, including GFAP+ and GFAP-populations is that they display reactivity, characterized by upregulation of GFAP, in response to injury (Bardehle et al., 2013). Here, we report that loss of *Lhx2* results in astrocytes developing some aberrant molecular signatures of reactivity in the absence of injury.

Our findings add unique astrocyte-specific roles to the key functions LHX2 plays in the development of the nervous system.

## Results

In the mouse, astrocyte production begins as neurogenesis ends, from embryonic day (E) 15.5, and peaks postnatally (Miller & Gauthier, 2007). LHX2 was identified as a regulator of the neuron-glia cell fate switch at embryonic stages (Subramanian et al., 2011). This transcription factor continues to be present in SOX2^+^ progenitors in the ventricular zone (VZ) and ALDH1L1^+^ cortical astrocytes (Figure 1A-D). We analyzed transcriptomic datasets from populations of purified cell types in the brain (Zhang et al., 2014) and discovered *Lhx2* expression to be high in mouse astrocytes compared with neurons, oligodendrocytes and microglia, at 1 month (Figure 1E).

**Figure 1:**
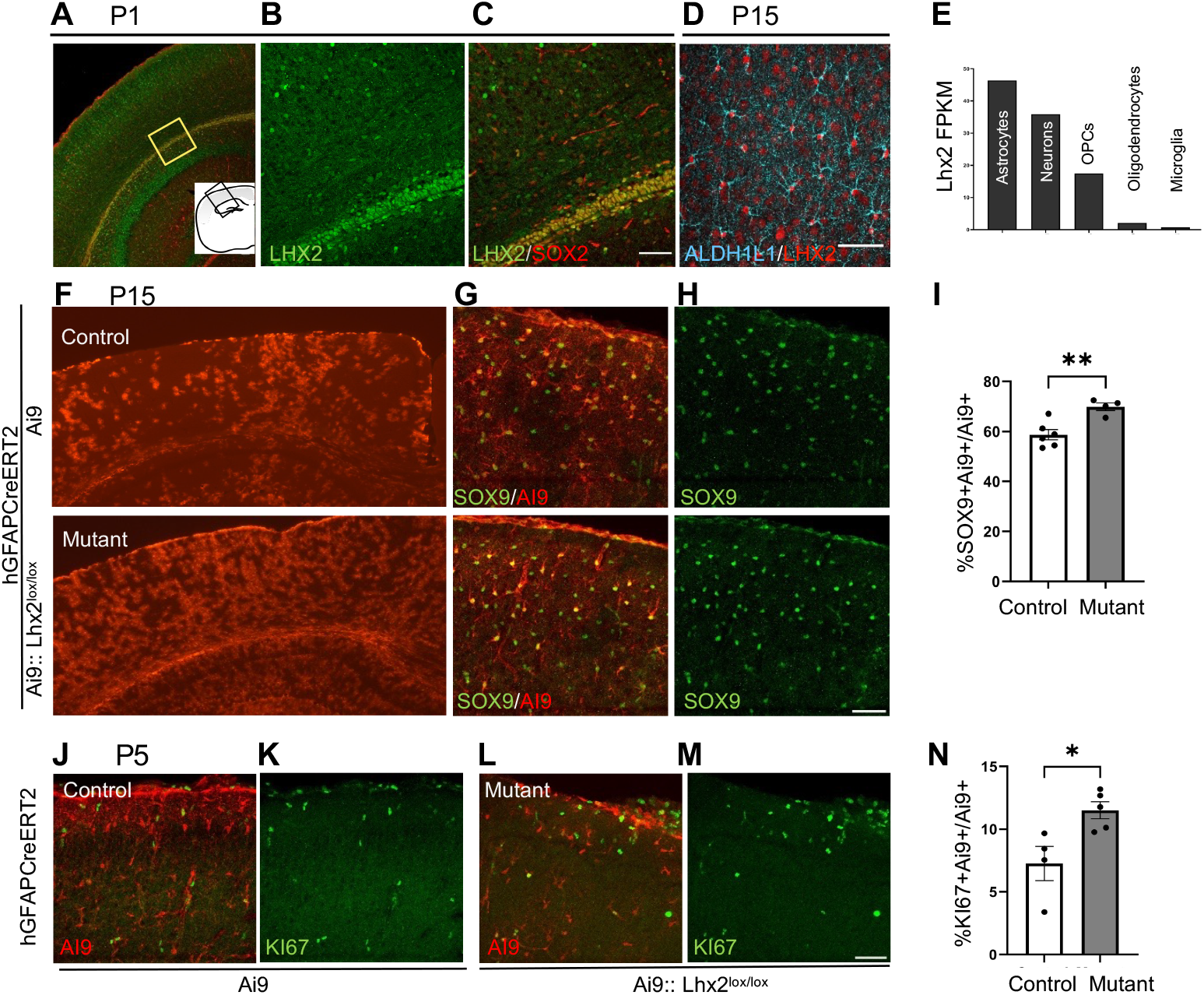
LHX2 loss in early postnatal cortex promotes astrogliogenesis via increase in proliferation. (A, B, C) LHX2 and SOX2 immunostaining in P1 progenitors. (B, C) are high magnification images of the ventricular zone of the somatosensory cortex (box in A). (D) LHX2 immunostaining in ALDH1L1+ astrocytes in the P15 cortex. (E) *Lhx2* expression in mouse brain (data analyzed from Zhang et al 2014). (F-I) P15 brains from hGFAPCreERT2:: Ai9 (control) and hGFAPCreERT2:: Ai9::Lhx2^lox/lox^(mutant) animals showing Ai9+ cells (F) co-labeled with SOX9 (G) or SOX9 alone (H). (I) Quantifications of the fraction of SOX9+ Ai9+ /Ai9+ cells: Control: 59%± 2; n = 3072 cells; N=6. Mutant: 70%± 1.5; n= 2282 cells; N= 4. Each N corresponds to a biologically independent replicate (individual mice). Statistical test: Unpaired t test with Welch’s correction, p=0.0024. (J-N) P5 brains stained for Ki67 show increased proliferation of Ai9+ cells in mutant (L, M) brains compared with controls (J, K). (N) Quantification of KI67+Ai9+/Ai9+ cells: Control: 7%± 1.36; n= 1436; N=4. Mutant: 12%±0.67; n=1876; N=5. Unpaired t test with Welch’s correction, p = 0.04. Error bars depict SEM. All scale bars are 50μm.

To examine the role LHX2 plays in astrocytes and their progenitors, we used a floxed *Lhx2* mouse line (Mangale et al., 2008) together with an *hGFAPCreERT2* driver (Ganat et al., 2006) to achieve conditional loss of *Lhx2*. The animals also carried an Ai9 reporter inserted in the ROSA 26 locus (Madisen et al., 2010) that allowed identification of cells that experienced Cre-driven recombination. Control animals carried *hGFAPCreERT2* together with the Ai9 reporter but did not carry the floxed *Lhx2* alleles. Since the *hGFAP* promoter drives expression both in progenitors that give rise to astrocytes, as well as in differentiated astrocytes, Ai9 fluorescence permits easy identification and scoring of astrocytes at later stages. Tamoxifen was administered at postnatal day (P) 1 and the brains were analyzed at different postnatal stages. At P3 and P5, Ai9+ cells display weak to undetectable LHX2 immunostaining upon loss of *Lhx2*, compared with control brains (Supplementary Figure S1 A-D).

### Loss of Lhx2 results in increased astrocyte proliferation

Upon tamoxifen administration at P1 and examination at P15, the Ai9+SOX9+ cells increased from 59% in controls to 70% upon loss of *Lhx2* (Figure 1F-I). We examined if this was due to an increase in proliferation using immunostaining for Ki67 at P5, a few days after tamoxifen administration (Clavreul et al., 2019; Ge, Miyawaki, Gage, Jan, & Jan, 2012). At this stage, the fraction of Ai9+Ki67+ colocalized cells increased from 7% in controls to 12% upon loss of *Lhx2*, indicating that enhanced proliferation is underway a few days after tamoxifen administration (Figure 1J-N).

### Cortical astrocytes display distinct Upper Layer (UL) versus Deep Layer (DL) molecular identities in the first postnatal week

Cortical astrocytes display distinct transcriptomic signatures depending on their localization in the UL or DL of the cortex at P15 (Bayraktar et al., 2020). We tested whether these transcriptomic differences are present earlier, during the first postnatal week. We mined the single cell (sc) RNA seq dataset from P4 somatosensory cortex (Di Bella et al., 2021). A Principal Component Analysis followed by an unsupervised Louvain clustering algorithm at a resolution of 0.5 split the cells into clusters (Supplementary Figure S1F) two of which expressed typical pan-astrocyte markers *Slc1a3, Aldoc, Aldh1L1* and *Gfap*. (Supplementary Figure S1G). Querying for established astrocytic layer-specific markers (Lanjakornsiripan et al., 2018; Bayraktar et al., 2020; Zhou et al., 2023) suggested that these two clusters correspond to the ULL (UL-like) and DLL (DL-like) respectively (Figure 2A, B, C and Supplementary Figure S1G, H, I). 1717 genes were differentially expressed between the two clusters, giving us a comprehensive tool to test for these two sub-categories (Supplementary Table S2).

**Figure 2:**
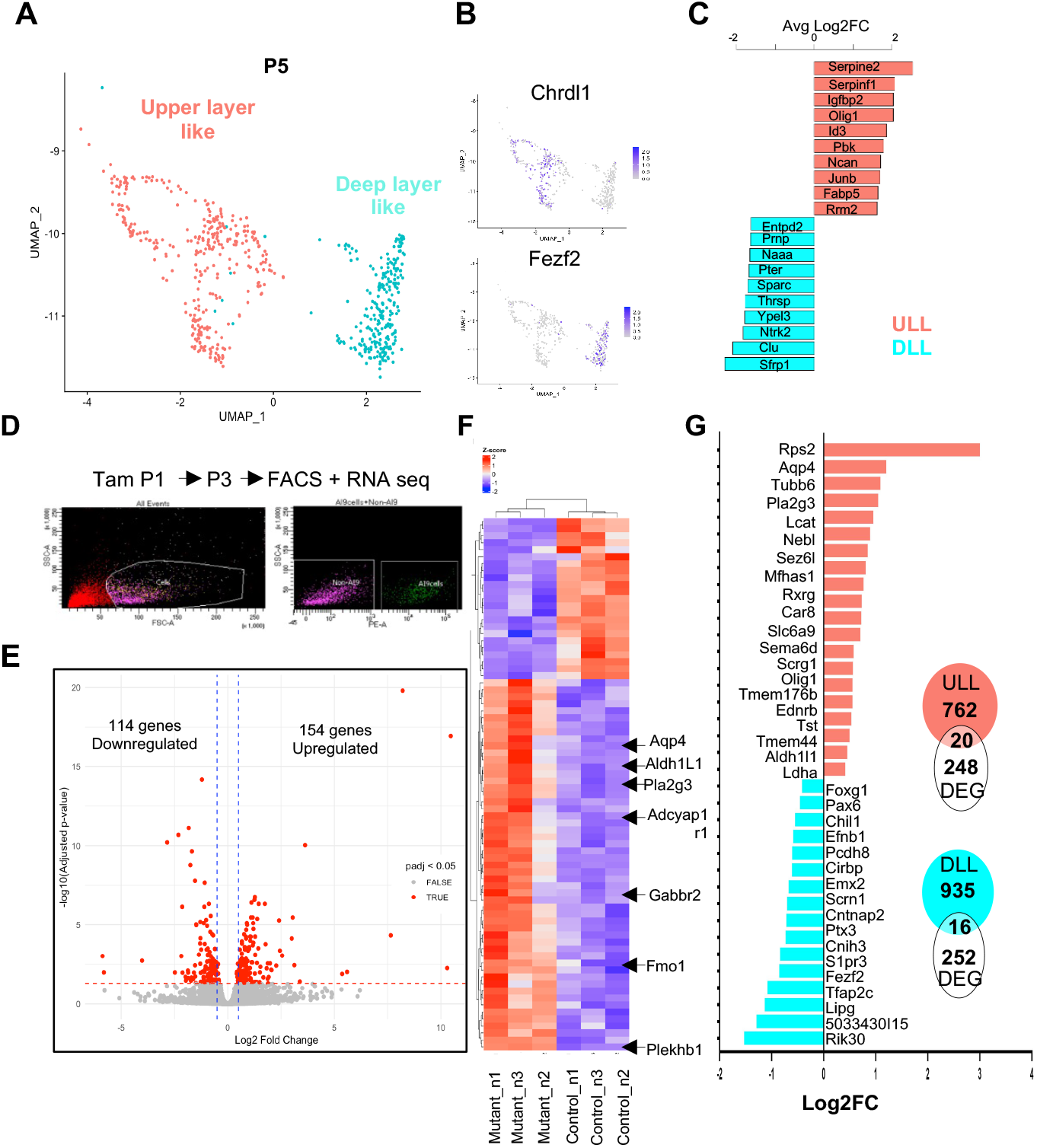
LHX2 loss results in increased UL astrocyte signatures. (A) A UMAP showing UL-like (ULL, salmon) and DL-like (DLL, cyan) astrocyte clusters analyzed from P4 single cell (sc) RNA seq data (Di Bella et al 2021). (B) Feature plots displaying the expression of one example of an UL- (*Chrdl1*) and a DL- (*Fezf2*) enriched candidate gene. (C) Average log2 Fold change of the top 10 differentially expressed genes between the two clusters (D) Workflow of the FACS based separation of P3 astrocytes followed by bulk RNA sequencing (E) Differentially expressed genes upon loss of *Lhx2* (F) Heatmap of the top 35 differentially expressed genes (DEG) between control and *Lhx2* mutant astrocytes (scale: red (high), blue (low) (G). Bar plot showing 20 upregulated ULL and 16 downregulated DLL genes upon Lhx2 loss. Inset, overlap of ULL and DLL genes in (A) with the DEGs in (E).

We used unsupervised Louvain clustering of E15.5 and E18.5 datasets (Di Bella et al., 2021) using the lowest resolution that creates two clusters. We found these clusters to display ULL and DLL signatures (Supplementary Figure S2). This indicates that astrocyte layer identity may be specified at the time of their generation, in progenitors or glial precursors.

### Loss of *Lhx2* results in differential dysregulation of UL-like versus DL-like astrocyte markers

We sought to examine whether LHX2 had a uniform role in the two spatially and molecularly distinct groups of UL and DL astrocytes. We used the same paradigm, *hGFAPCreERT2 :: Ai9* for controls, and *hGFAPCreERT2 :: Ai9 :: Lhx2* ^*lox/lox*^, administered tamoxifen at P1, isolated Ai9+ cells using FACS at P3, and performed RNAseq (Figure 2D). This age was selected to allow sufficient time for transcriptomic dysregulation upon loss of *Lhx2*. The sequencing reads were examined for excision of *Lhx2* exons 2 and 3, which are floxed in the conditional *Lhx2* allele (Mangale et al., 2008). As expected, control samples contained reads for all the exons of the *Lhx2* gene while the mutant samples lacked the exon 2 and 3 reads showing that the knockout was successful (Supplementary Figure S1E). We performed differential gene expression analysis using DESeq2 between control and *Lhx2* mutant samples and it revealed 114 downregulated genes and 154 upregulated genes (Figure 2E, Supplementary Table S1). Hierarchical clustering showed that control and mutant replicates clustered together (Figure 2F). There was an upregulation of pan-astrocytic markers *Aldh1L1* and *Aqp4* as well as several astrocyte enriched ligands and receptors such as *Pla2g3, Adcyap1r1, Gabbr2, Pla2g7, Ednrb*. Interestingly, *Slc25a34* and *Fmo1*, which are enriched in UL astrocytes, (Bayraktar et al., 2020; Lanjakornsiripan et al., 2018) were also upregulated (Figure 2E, F). To examine if upper layer astrocyte signatures were altered upon loss of *Lhx2*, we compared the genes dysregulated in *Lhx2* mutant astrocytes with the 1717 genes that we identified as differentially expressed in the ULL and DLL P4 astrocyte clusters (Figure 2G; Supplementary Table S2). Of these, 20 genes associated with ULL astrocytes were upregulated and 3 were downregulated, whereas 16 genes associated with DLL astrocytes were downregulated and 5 were upregulated upon loss of *Lhx2 (*Figure 2G, Supplementary Figure S3A-E).

These findings suggested that LHX2 may not play a uniform role in astrocytes occupying different cortical layers. Therefore, we examined whether the increased proliferation of astrocytes seen upon loss of *Lhx2* was uniform across the cortical UL and DL.

### UL astrocytes divide more than DL astrocytes upon loss of *Lhx2*

First, we examined *Ki67* expression from E15.5 - P4 in the scRNAseq dataset of (Di Bella et al., 2021), and found it to be higher in UL than in DL cells. These are likely to include not only astrocytes, but also their progenitors or glial precursors, present at different developmental stages (Figure 3A, Supplementary Figure S2C, F). Therefore, the higher proliferation of UL astrocytes reported in the first postnatal week (Clavreul et al., 2019) appears from the earliest stages of astrocyte genesis in the embryo.

**Figure 3:**
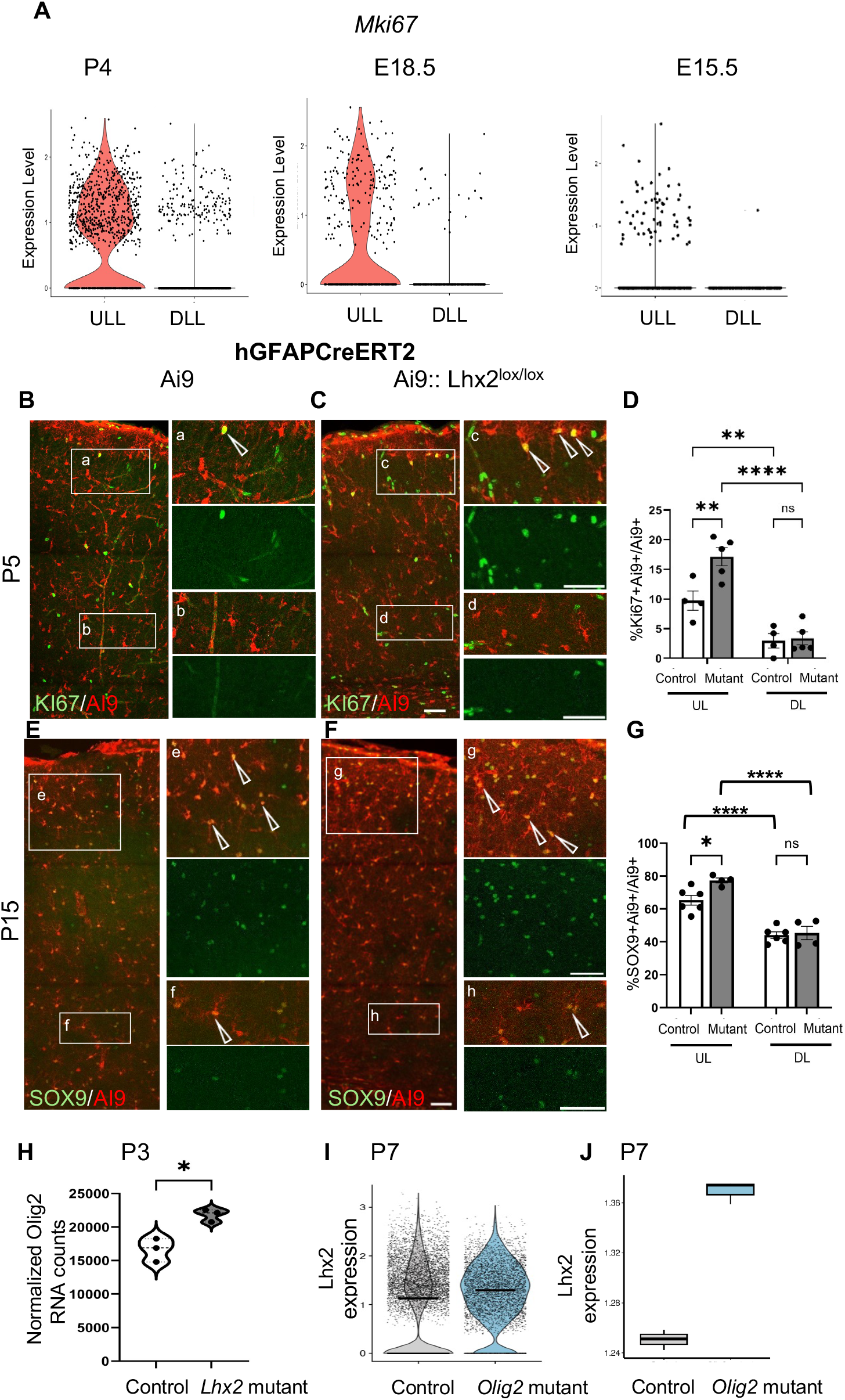
Differential proliferation of UL and DL astrocytes. (A) Violin plots showing *mKi67* expression in E15.5, E18.5 and P4 UL-like (ULL) versus DL-like (DLL) astrocytes/ glial precursors/progenitors (dataset from Di Bella et al. 2021). (B-D) At P5, KI67 immunostaining (B, C) and quantification (D) shows increased proliferation of Ai9+ cells in the control UL (10% ± 1.61) versus the DL (3%± 1.2, p=0.0054); this is further increased in the mutant UL (17%± 1.52, p=0.002) but not in the DL (3%± 1.07, p= 0.8). Astrocytes in the white matter were not included in the DL counts. Statistical test: Two-way ANOVA with Fisher’s LSD multiple comparison test. Control: n= 1436 Ai9 cells; N= 4. Mutant: n= 1876 cells; N= 5 independent biological replicates. (E-G) At P15, SOX9 immunostaining (E, F) and quantification (G) shows an increase in the SOX9+Ai9+ fraction of Ai9+ cells in the control UL (65% ± 2.96) versus the DL (44% ± 1.93, p <0.0001); this is further increased in the mutant UL (77% ± 1.52, p= 0.03) but not in the DL (45% ± 4.01, p =0.98). Two-way ANOVA with Sidak’s multiple comparison test). Control: n= 3054 Ai9 cells; N= 6. Mutant: n= 2221 cells; N= 4 independent biological replicates. Regions in white boxes in (B, C, E, F) marked (a-h) are presented at high magnification in adjacent panels showing the red-green overlay and a single green channel. Open arrowheads mark double labelled cells. (H) *Olig2* expression in Ai9+ cells is increased upon loss of *Lhx2*. Data from normalised counts obtained from RNA seq (Figure 2D) (p=0.01, Unpaired t test with Welch’s correction). Error bars represent SEM. (I, J) scRNAseq from P7 astrocytes (Zhou et al., 2023), reveals increased *Lhx2* expression upon loss of *Olig2* displayed in a violin plot (I) and pseudobulk analysis of the same data displayed in a box-whisker plot (J; Unpaired t test, p= 7.246e-05) Panels in B,C,E,F and K are composites produced from multiple images using the multi area acquisition feature on the confocal FLUOVIEW^TM^ FV 3000 software. All scale bars are 50µm

KI67 immunostaining at P5 in control sections was consistent with these findings, i.e., the UL contained a greater percentage of the Ai9+cells that were also Ki67+ than the DL (Figure 3B). Notably, we found that this difference was further enhanced upon loss of *Lhx2*. The Ai9+Ki67+ fraction of the total Ai9+ cells for the mutant UL increased significantly compared to control UL, whereas the corresponding counts for the mutant DL remained similar to control DL (Figure 3B-D). We also examined the astrocyte marker SOX9 at P15 when local astrocytic proliferation is nearly complete. Ai9+SOX9+ cells were scored based on their presence in UL versus DL in sections from the same experiment shown in Figure 1I, that showed an overall increase in SOX9+ cells upon loss of *Lhx2*. We observed an increase in Ai9+SOX9+ fraction in the mutant UL, from 65% in controls to 77% in the mutant UL, but no change in the DL between the two groups (Figure 3E, F, G). Together, the data indicate that loss of *Lhx2* has a differential effect on astrocyte proliferation and explains why UL astrocyte signatures increased in the transcriptomic analysis (Figure 2G).

### *Lhx2* and *Olig2* regulate aspects of astrocyte development in the UL and DL respectively

Regulatory mechanisms that determine astrocyte numbers are not well understood. Loss of *Olig2* in astrocytes has been reported to produce a molecular identity transition from grey matter to white matter (WM) astrocytes (Zhou et al., 2023). Since WM astrocytes share several molecular features with those in the DL (Bayraktar et al., 2020), this suggests that OLIG2 may regulate aspects of UL-like identity. *Olig2* expression increases upon loss of *Lhx2*, consistent with an increase in UL-like astrocytes (Figure 3H).

We analyzed the scRNAseq data from P7 astrocytes in Zhou et al., (2023) and found an increase in *Lhx2* expression upon loss of *Olig2* (Figure 3I, J), indicating that OLIG2 suppresses *Lhx2* in astrocytes. These data suggest a possible Olig2-Lhx2 mediated regulation of astrocyte proliferation.

hGFAP Cre labels both astrocyte and oligodendrocyte precursor cells (OPCs; (Weng et al., 2019). We examined other OPC markers, *Sox10, Pdgfra, Olig1, Epn2* and *Zfp36l1* in the P3 RNA seq data from Ai9+cells, and found that loss of *Lhx2* results in increased expression of each of these genes (Supplementary Figure S4A). However, OLIG2 immunostaining at P5 does not reveal a difference in the fraction of OLIG2+ Ai9+/ Ai9+ cells upon loss of *Lhx2* (Supplementary Figure S4B). Since oligogenesis reaches its peak later than astrogliogenesis, further analysis at suitable stages, using a Cre driver specific to the OPC lineage, would clarify the role of LHX2 in this distinct glial lineage.

### Long-term consequences of loss of *Lhx2* in astrocytes

In the retina, *Lhx2* mutant Muller glia upregulate GFAP expression 30 days after tamoxifen-driven induction of Cre recombination (de Melo et al., 2012), similar to what is seen in reactive astrocytes in response to injury (Sofroniew, 2005). We examined cortical astrocytes arising from loss of *Lhx2* at P1 for typical markers of reactive gliosis, GFAP and VIMENTIN (Buffo et al., 2008).

At 3 months, GFAP and VIMENTIN, is seen in pial astrocytes in the marginal zone (MZ) and those in and near the white matter (Figure 4A, C). VIMENTIN is also present in endothelial cells. In *Lhx2* mutants, both these markers were seen in grey matter astrocytes across the radial extent of the cortex. These astrocytes were also Ai9+ indicating that they arose from cells that experienced loss of *Lhx2* from P1 (Figure 4A, C). The fraction of Ai9+ astrocytes that were also VIMENTIN+ increased from 19% and 2% in the control UL and DL respectively, to 61% and 42% in the mutant UL and DL respectively (Figure 4B). An examination of the stack (Supplementary Figure S5A, Movie 1 and 2) reveals enhanced VIMENTIN signal in *Lhx2* mutants. Similarly, the fraction of Ai9+ astrocytes that were also GFAP+ increased from 10% and 19% in the control UL and DL respectively, to 47% and 48% in the mutant UL and DL respectively (Figure 4D).

**Figure 4:**
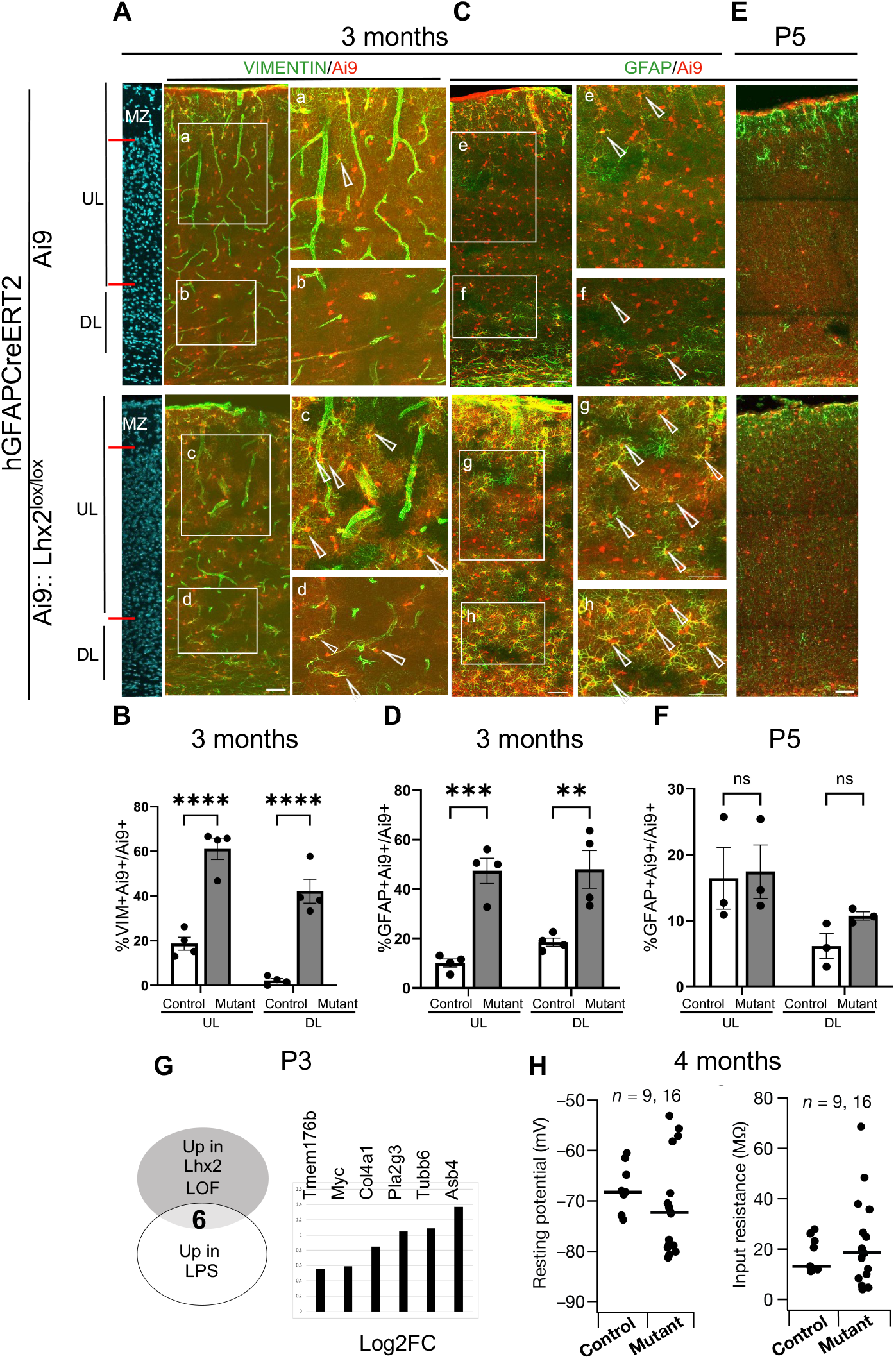
Long-term consequences of LHX2 loss in astrocytes. (A-D) Sections of the cortex at 3 months stained with DAPI (blue, A) to identify the marginal zone (MZ), UL and DL. GFAP immunostaining labels astrocytes(C) and VIMENTIN immunostaining also labels endothelial cells (A). VIMENTIN+ (A) or GFAP+ (C) astrocytes are seen across the radial extent of the cortex in mutants (open arrowheads, A, C), but not in controls. Regions in white boxes in (A, C) marked (a-h) are presented at high magnification in adjacent panels. (B) Quantifications of VIMENTIN+Ai9+/ Ai9+ cells: Control UL: 19%± 2.94, Mutant UL: 61%±4.78, p < 0.0001; Control DL: 2%± 0.92 Mutant DL: 42%± 5.3, p< 0.0001). Control: n = 1326, Mutant:n = 1079, N=4 (biologically independent replicates). (D) Quantification of GFAP+Ai9+/ Ai9+ cells: Control UL: 10%± 1.6, Mutant UL: 47%± 5, p=0.0002; Control DL: 19%± 1.6 Mutant DL: 48%± 7.6, p= 0.0017). Control: n = 2640, Mutant: n= 1975, N=4 (biologically independent replicates) (E) GFAP Immunostaining at P5. (F) Quantification of GFAP+Ai9+/ Ai9+ cells: Control UL: 16%± 4.7, Mutant UL: 17%± 4, p=0.9; Control DL: 6%± 1.9 Mutant DL: 11%± 0.61, p= 0.6. (Control: n = 1412, Mutant: n= 778, N=3). For D-F, a two-way ANOVA test with Sidak’s multiple comparison test was used. Error bars depict SEM. (G) Genes upregulated upon *Lhx2* LOF in astrocytes at P3 (this study) overlapped with LPS-induced reactive astrocyte signatures (Hasel et al. 2021). 6 genes common to both datasets with their Log2Fold change upon *Lhx2* LOF. (H) Beeswarm plots of resting membrane potential (*V*_RMP_) and input resistance (*R*_in_) obtained from control and mutant LII/III cortex astrocytes using whole cell patch clamp configuration. The thin lines indicate the respective median values. *p*=0.19 for *V*_RMP_ and *p*=0.93 for *R*_in_ (Wilcoxon rank sum test). All scale bars are 50µm. Panels in A-C are composites each produced from multiple images using a multi area acquisition setting on the confocal FLUOVIEW^TM^ FV 3000 software.

We examined whether the upregulation of GFAP manifested earlier, at P5. We found GFAP staining to appear similar in control and mutant brains, with no change in the number of Ai9+ GFAP+ astrocytes in *Lhx2* mutant brains (Figure 4E, F). However, some genes that were upregulated upon loss of *Lhx2* in the P3 RNAseq were identified as markers of reactive astrocytes in an LPS-induced model (Figure 4G; (Hasel, Rose, Sadick, Kim, & Liddelow, 2021)). This suggests that although GFAP itself does not appear upregulated upon loss of *Lhx2* by P5, aspects of a reactive gliosis-like state may be underway at this stage. Examination of microglial marker IBA1 reveals no increase in the number of microglia, consistent with the absence of a general inflammatory state (Supplementary Figure S5B).

The observation that reactive-gliosis-like signatures arise in the absence of injury suggests that LHX2 may participate in mechanisms involving suppressing aberrant reactive signatures in astrocytes. Injury-driven mechanisms may act via suppression of LHX2 to upregulate these signatures. Since OLIG2 is known to regulate aspects of injury-related reactive gliosis, including astrocyte proliferation, (Chen et al., 2008), one intriguing line of exploration is a potential link between LHX2 and OLIG2 in this context. An alternate possibility is that loss of *Lhx2* results in increased proliferation of the GFAP+ pial astrocytes population which may migrate to occupy the radial extent of the cortex. Further investigation is needed to examine how LHX2 modulates astrocyte properties including molecular features in maturity.

We tested whether the electrophysiological properties of the astrocytes, produced as a result of loss of *Lhx2* at P1, were altered in maturity (> 3 months; Figure 4H; Supplementary Figure S6). First, the non-neuronal nature of the cell being patched was confirmed by ascertaining that no action potentials were elicited even for large current injections up to 1250 pA. The resting membrane potential (Figure 4H), input resistance (Figure 4H, Supplementary Figure S6A), Sag (Supplementary Figure S6B-C), and temporal summation (Supplementary Figure S6D-E) of control and mutant astrocytes were not significantly different from each other. We characterized the frequency-dependent properties of these cells with the chirp stimulus (Supplementary Figure S6F;(Ashhad & Narayanan, 2016; Mishra & Narayanan, 2020, 2022). We found no pronounced difference in the gain (Supplementary Figure S6G) or the impedance phase (Supplementary Figure S6K) of the astrocytes through the measured frequency range of 0–15 Hz. Maximal impedance amplitude (Supplementary Figure S6H), resonance frequency (Supplementary Figure S6I), resonance strength (Supplementary Figure S6J), and total inductive phase (Supplementary Figure S6L) of control and mutant astrocytes were not significantly different from each other. Together, these measurements revealed that the basic electrophysiological properties of cortical astrocytes were comparable to controls despite the loss of *Lhx2*.

In mice and humans, cortical astrocytes are produced from progenitors residing in the ventricular and subventricular zones (VZ and SVZ respectively), and also via self-amplification, i.e. local proliferation in the cortex (D. E. Allen et al., 2022; Clavreul et al., 2019; Ge et al., 2012). Given the regional, molecular and morphological heterogeneity in the astrocyte population, it is imperative to understand the mechanisms that regulate astrogliogenesis of these different sub-groups. Transcription factor LHX2 is detected in progenitors and in astrocytes at all stages until adulthood. In wild type brains, astrocytes residing in the cortical UL normally proliferate more than those in the DL (Figure 3A, (Clavreul et al., 2019)). We provide evidence that LHX2 specifically suppresses UL, not DL astrocyte division in the early postnatal window, suggesting a mechanism to restrict unrestrained UL astrocytic proliferation. This is the first report of a layer-specific regulator of astrocyte proliferation.

Our analysis of the single cell transcriptomic data (Di Bella et al., 2021) revealed separate clusters of cells that display ULL and DLL signatures as early as E15.5 when progenitors are in the early phase of astrogliogenesis (Supplementary Figure S2). This indicates that astrocyte layer specification occurs from the start of their genesis, and opens a new avenue of exploration in the acquisition of layer identity of astrocytes, similar to what has been explored for neuronal sub-types for over three decades. Indeed, a recent study identified bHLH transcription factor OLIG2 to be a suppressor of DL astrocyte fate (Zhou et al., 2023). Loss of *Olig2* also causes upregulation of *Lhx2* expression. This suggests that LHX2 and OLIG2 may work in a complementary manner to regulate the numbers of cortical astrocytes, LHX2 by suppressing overproliferation in the UL, and OLIG2 by suppressing fate specification of DL astrocytes.

In summary, this study uncovers a novel role for LHX2 in the control of cortical layer-specific astrocyte proliferation and also implicates this factor in regulating the reactive response of astrocytes that is normally seen in the context of injury.

## Acknowledgements

We thank our animal house in-charge, Shital Suryavanshi and the animal house staff of the Tata Institute of Fundamental Research, Mumbai (TIFR) for their excellent support. We thank Deepak Modi, Vainav Patel and Sameen Khan for generous support and access to the FACS Facility at National Institute of Reproductive and Child Health (NIRRCH), Mumbai. We also thank Hema Bagul at the TIFR FACS facility for her assistance. We thank Radhika Dharap for her assistance in formatting the graphical representations in Figures 3 and 4.

## Competing Interests

The authors declare no financial or competing interests.

## Funding

This work was funded by the DST INSPIRE Faculty Fellowship awarded to AI (Faculty Registration no: IFA18-LSBM210), the Department of Atomic Energy (DAE), Govt. of India (Project Identification no. RTI4003, DAE OM no. 1303/2/2019/R&D-II/DAE/2079)

## Data availability

The Fastq files of the RNA seq data generated from P3 sorted astrocytes for control and Lhx2 mutants have been uploaded as an SRA submission with the accession ID: PRJNA114957

## Materials and Methods

### Ethics statement

All animal protocols were approved by the Institutional Animal Ethics Committee of the Tata Institute of Fundamental Research (TIFR-IAEC). All the electrophysiology experiments mentioned in this study were approved by the Institute Animal Ethics Committee of the Indian Institute of Science, Bangalore. Both of these committees ensure that all animal studies were conducted in accordance with ethical guidelines.

### Mice

The *Lhx2l*^*lox/lox*^ mouse line used in this study has been previously described in Mangale et al 2008. *hGFAPCreERT2* (Strain #:012849), and *Ai9* reporter mouse line (Strain #:007909) are purchased from the Jackson laboratory.

The *Lhx2* floxed allele (Mangale et al., 2008) was generated using a targeting construct designed to recapitulate the *Lhx2* null animal (Porter et al., 1997). This mutant which was created by targeted disruption of exons 2 and 3 of *Lhx2* that encode the LIM domains and the domain linking the LIM domains and homeodomain of Lhx2. Splicing between exons one and four results in a frameshift such that no homeodomain containing peptide can be produced (Porter et al., 1997). Only exon 1 is in-frame, which encodes neither the LIM domains nor the HD. Immunostaining for LHX2 in *hGFAPCreERT2 :: Ai9 :: Lhx2l*^*lox/lox*^ brains shows no detectable LHX2 protein in the Ai9+ astrocytes (Supplementary Figure S1A-C).

Mice homozygous for floxed alleles of *Lhx2* ^*lox/lox*^ *and Ai9 reporter (Lhx2* ^*lox/lox*^*;Ai9/Ai9)* were crossed with mice carrying Cre recombinase driven by the human GFAP promoter (*hGFAPCreERT2/+; Lhx2* ^*lox/lox*^) to generate *Lhx2* conditional knockout (cKO) (*hGFAPCreERT2/+; Lhx2* ^*lox/lox*^*;Ai9/Ai9)*. Controls were *hGFAPCreERT2* animals crossed to *Ai9/Ai9* animals. Animals of either sex were used in the study. All animals were kept at an ambient temperature and humidity, with a 12h light-dark cycle and food available ad libitum. Noon of the day of the vaginal plug was designated as embryonic day 0.5 (E0.5).

Neonatal pups were genotyped by nail and tail clip method. In this procedure, the tip of the nail was clipped from the digits 3 and 4 in various combinations from the 4 paws to identify pups from each other and the tail clip was taken to genotype.

Primers used for genotyping were:

Detection of Cre for *hGFAPCreERT2*

Cre F: 5′ATTTGCCTGCATTACCGGTC3′,

Cre R: 5′ATCAACGTTTTCTTTTCGG3′ . Cre-positive DNA shows a band at 350 bp.

*Lhx2* ^*lox/lox*^ F: 5’ACCGGTGGAGGAAGACTTTT3’,

*Lhx2* ^*lox/lox*^ R: 5’CAGCGGTTAAGTATTGGGACA3’. The band sizes for this PCR are as follows:

Wild-type: 144 bp, *Lhx2* floxed allele: 188 bp.

### Tamoxifen administration

Tamoxifen (Sigma, T5648) was prepared in corn oil (Sigma, C82687) with a concentration of 20 mg/ml and dissolved overnight at 37°C on a shaker. Cre positive pups and the dam were administered Tamoxifen at 40 mg/kg at P1.

### Tissue Preparation

Neonatal pups were euthanized on ice and brains were harvested in cold Phosphate Buffer Saline and fixed overnight in 4% (wt/vol) paraformaldehyde. Adult animals were administered Thiosol (0.5mg/ml) at 0.04mg/kg as an intraperitoneal injection to be anesthetized and transcardially perfused with 4% (wt/vol) paraformaldehyde in phosphate buffer, followed by overnight fixation. Prior to sectioning, they were transferred to 30% (wt/vol) sucrose-PBS for cryoprotection during the sectioning process. The brains were sectioned at 30μm and 40μm using a freezing microtome (Leica SM2000R). In all of the analysis (except electrophysiology), the primary somatosensory cortex (S1) was analysed.

### Immunofluorescence

Brains were sectioned (30 or 40 μm) and sections from P15 and adult brains were processed in a 4 well plate for immunofluorescence. Sections from P5 brains were mounted on Superfrost plus glass microscope slides (Cat no 71869-10), and dried for 3 h at 37°C. The rest of the procedure was common to all ages. Sections were hydrated in a phosphate buffer by washing the sections thrice for 5 mins each. Antigen retrieval was performed at 90°C–95°C in 10 mM sodium citrate buffer (pH 6.0) for 10 mins. Following this step, sections were subjected to a blocking solution comprising 5% (v/v) horse serum in phosphate buffer with 0.3% (v/v) Triton X-100 (0.1% for P5) (Sigma; X100) for 1 h at room temperature. Incubation with primary antibody was performed in a phosphate buffer containing 0.3% or 0.1% (v/v) Triton X-100 and 2.5% (vol/vol) horse serum at 4°C overnight. The following day, sections were washed in phosphate buffer thrice for 5 mins each, followed by the appropriate secondary antibody (prepared in phosphate buffer containing 0.3% or 0.1% (v/v) Triton X-100) for two hours at room temperature. This was followed by three washes for 5 minutes each in phosphate buffer and DAPI (Molecular Probes, Cat no D1306) staining for 10 minutes, after which the sections were washed with phosphate buffer thrice (5 mins each). The slides were then mounted with Fluoroshield (Sigma, Cat no F6057 or F6182). Information on antibodies and their catalog numbers are mentioned in SI Appendix.

### Image acquisition and analysis

Images were acquired with a FV1200 or FV3000 Olympus inverted confocal microscope equipped with an oil immersion objective lens (X40) or (X10). Confocal images were processed with the Fiji package of ImageJ (NIH).

### Quantification of SOX9, KI67 and VIMENTIN

Sections immunostained for each marker together with RFP (for Ai9 reporter) were imaged on the FV3000 Olympus inverted confocal microscope equipped with an oil immersion objective lens(X40). Images for each experiment were acquired under similar PMT settings in sequential scan mode.

The cortical layers were identified by DAPI staining and sections were manually scored within the UL or DL for the number of Marker+ Ai9+ fraction of the total Ai9+ cells. Since this analysis was restricted to UL and DL, white matter astrocytes were not included.

### Patch-clamp electrophysiology experiments

1. (2 males and 3 females) mice belonging to the control group between 4–7 months old and 6 (3 males and 3 females) Lhx2 mutant mice between 3–7 months old were used for in vitro patch-clamp electrophysiology experiments.

### Slice preparation for in vitro patch-clamp recording

Mice were anesthetized by intraperitoneal injection of a ketamine-xylazine mixture. After onset of deep anesthesia, assessed by cessation of toe-pinch reflex, transcardial perfusion of ice-cold cutting solution was performed. The cutting solution contained (in mM) 2.5 KCl, 1.25 NaH_2_PO_4_, 25 NaHCO_3_, 0.5 CaCl_2_, 7 MgCl_2_, 7 dextrose, 3 sodium pyruvate, and 200 sucrose (pH 7.3, ∼300 mOsm) saturated with carbogen (95% O_2_, 5% CO_2_). Thereafter, the brain was removed quickly and 350-µm thick near-horizontal slices were prepared with a vibrating blade microtome (Leica Vibratome) while submerged in ice-cold cutting solution saturated with carbogen. The slices were then incubated for 10–15 min at 34° C in a holding chamber containing a holding solution (pH 7.3, ∼300 mOsm) with the composition of (in mM) 125 NaCl, 2.5 KCl, 1.25 NaH_2_PO_4_, 25 NaHCO_3_, 2 CaCl_2_, 2 MgCl_2_, 10 dextrose, and 3 sodium pyruvate saturated with carbogen. Thereafter, the slices were kept in the holding chamber at room temperature for at least 30 min before recordings started.

### Whole-cell current-clamp recordings

For electrophysiological recordings, slices were transferred to the recording chamber and were continuously perfused with carbogenated artificial cerebrospinal fluid (ACSF-extracellular recording solution) at a flow rate of 2–3 mL/min. All astrocytic recordings were performed under current-clamp configuration at physiological temperatures (32–35°C) achieved through an inline heater that was part of a closed-loop temperature control system (Harvard Apparatus). The ACSF contained (in mM) 125 NaCl, 3 KCl, 1.25 NaH_2_PO_4_, 25 NaHCO_3_, 2 CaCl_2_, 1 MgCl_2_, and 10 dextrose (pH 7.3; ∼300 mOsm). Slices were first visualized under a 10× objective lens to visually locate layer II/III of the somatosensory cortex (S1). A 63× water-immersion objective lens was employed to perform visually guided patch-clamp recordings from mCherry-labeled astrocytes in superficial layers of cortex, through a Dodt contrast microscope (Carl Zeiss Axioexaminer). Whole cell current-clamp recordings were performed from layer II/III S1 astrocytes with a Dagan BVC-700A amplifier.

Borosilicate glass electrodes with electrode tip resistance between 4 and 8 MΩ were pulled (P-97 Flaming/Brown micropipette puller; Sutter) from thick glass capillaries (1.5-mm outer diameter and 0.86-mm inner diameter; Sutter) and used for patch-clamp recordings. The pipette solution contained (in mM) 120 K-gluconate, 20 KCl, 10 HEPES, 4 NaCl, 4 Mg-ATP, 0.3 Na-GTP, and 7 K_2_-phosphocreatine (pH 7.3 adjusted with KOH; osmolarity ∼300 mOsm). Series resistance was monitored and compensated online with the bridge-balance circuit of the amplifier. Experiments were discarded only if the initial resting membrane potential was more depolarized than –50 mV or if series resistance rose above 50 MΩ or if there were fluctuations in temperature and ACSF flow rate during the experiment. Voltages have not been corrected for the liquid junction potential, which was experimentally measured to be ∼8 mV. Astrocytic response to a 250-pA hyperpolarizing current pulse was continuously monitored to observe and correct series resistance changes using the bridge balance circuit throughout the course of the experiment.

### Pharmacological blockers

All recordings were performed in presence of synaptic receptor blockers in the ACSF. Drugs and their concentrations used in the experiments were 10 µM 6-cyano-7-nitroquinoxaline-2,3-dione (CNQX), an AMPA receptor blocker; 10 µM (+) bicuculline and 10 µM picrotoxin, both GABA_A_ receptor blockers, and 2 µM CGP55845, a GABA_B_ receptor blocker (all synaptic blockers from Abcam) in the ACSF.

### Electrophysiological measurements

We characterized S1 astrocytes with several electrophysiological measurements using standard protocols (Narayanan & Johnston, 2007); (Ashhad & Narayanan, 2016; Mishra & Narayanan, 2020, 2022; Narayanan & Johnston, 2008), detailed below. Resting membrane potential, *V*_RMP_ was measured as the voltage at which the cell rested when no current was injected (Figure 4H). Input resistance (*R*_in_) was measured as the slope of a linear fit to the steady-state voltage-current (*V–I*) plot obtained by injecting current pulses of amplitudes spanning –250 to 250 pA, in steps of 50 pA (Supplementary Figure S5A). Sag ratio was measured from the voltage response of the cell to a hyperpolarizing current pulse of 250 pA (Supplementary Figure S5B, C). Sag ratio was defined as (V _initial_/V_ss_), where V_ss_ and V _initial_ depict the steady-state and peak (during the initial 50-ms period after current injection) voltage deflections (from V_RMP_), respectively. To assess temporal summation, five alpha excitatory postsynaptic potentials (α-EPSPs) with 50-ms interval were injected as currents of the form *I*_*α*_ = *I*_*max*_ *t exp*(– *αt*), with *α* = 0.1 ms^−1^ . Temporal summation ratio (*S*_*α*_) in this train of five EPSPs (Supplementary Figure S5*D, E*) was computed as *E*_*last*_ /*E*_*first*_, where *E*_*last*_ and *E*_*first*_/, were the amplitudes of the last and first EPSPs in the train, respectively.

The chirp stimulus, a sinusoidal current with its frequency linearly spanning 0–15 Hz in 15 s and of constant amplitude was used for characterizing the impedance profiles (Supplementary Figure S5F-J). The voltage response of the astrocyte to the chirp current stimulus injection was recorded at *V*_RMP_. The ratio of the Fourier transform of the voltage response to the Fourier transform of the chirp stimulus formed the impedance profile. The frequency at which the impedance amplitude reached its maximum was the resonance frequency (*f*_R_). Resonance strength (*Q*) was measured as the ratio of the maximum impedance amplitude to the impedance amplitude at 0.5 Hz. Total inductive phase (ϕ_L_) was defined as the area under the inductive part of the impedance phase profile as a function of frequency (Supplementary Figure S5K-L). No action potential firing was observed when higher currents, in the range of 250–1250 pA in steps of 250 pA, were injected into the astrocytes.

### Analyses of electrophysiological data and statistics

All data acquisition and analyses were performed with custom-written software in IGOR Pro (WaveMetrics), and statistical analyses were performed using Wilcoxon rank sum test in R computing package (http://www.r-project.org/).

### Fluorescence Activated Cell Sorting

Control and *Lhx2* mutants were obtained as described above. Tamoxifen (40 mg/kg) was administered at P1 and brains were harvested at P3. The brains were dissected in cold Hank’s Balanced Salt Solution (Thermo Fisher Scientific, Cat no: 14170112), the meninges were removed and the cortical tissue was dissected out and collected in RNase free Eppendorf tubes. The tissue was spun down at 400 g for 1 min and the excess buffer was removed. The tissue was then subjected to trypsinization (0.25% trypsin prepared in HBSS without Calcium and Magnesium) for 10 mins at 37°C on a shaker post which it was spun down for 5 mins at 400g. The tissue was then neutralized with 10% FBS in HBSS (with Calcium and Magnesium) and using trituration a homogenous cell suspension was obtained. This suspension was spun down at 400g for 5 mins and the resulting cell pellet was resuspended in HBSS (with Calcium and Magnesium) to obtain a homogeneous suspension. The suspension was then passed through a 70 micron cell strainer to remove any debris. FACS was performed using BD FACSAria^TM^ Fusion (BD Biosciences) with the 568 laser using a 100μm nozzle. Singlets were selected using forward scatter and side scatter.

Cells were selected for collection based on their RFP signal. A small sample of cells were re-sorted to ensure the sorting efficiency during each collection round. For sequencing, 3 replicates of control and mutant condition each containing approximately 300000 cells were collected. Cells were collected directly in the Lysis buffer (Buffer RLT, Qiagen, Cat no 79216). For 350µl of the RLT buffer, 3.5µl Beta Mercaptoethanol was added, mixed well and frozen down (-80°C).

### RNA extraction and cDNA synthesis

RLT buffer (Buffer RLT, Qiagen, Cat no 79216) containing frozen sorted cells for control and Lhx2 mutant were sent to Medgenome Labs Ltd (Bangalore) where RNA extraction, cDNA synthesis and sequencing was performed. RNA was extracted from cells frozen in the RLT buffer using the RNeasy mini kit (Cat no 74104). The Extracted RNA samples were quantified using Qubit RNA BR Assay (Invitrogen, Cat no Q10211) / Qubit RNA Assay HS (Invitrogen, Cat no Q32852). RNA purity was checked using QIAxpert and RNA integrity was checked on RNA screen tape (Agilent, Cat no 5067-5576).

NEB Ultra II directional RNA-Seq Library Prep kit (Cat no E7760L) protocol was used to prepare libraries for mRNA sequencing. An initial Concentration range of 100-500 ng of total RNA was taken for the assay. mRNA molecules were captured using magnetic Poly(T) beads (NEB, Cat no E7490L). Following purification, the enriched mRNA pool were fragmented using divalent cations under elevated temperatures. The cleaved RNA fragments were copied into first-strand cDNA using reverse transcriptase. Second strand cDNA synthesis was performed using DNA Polymerase I and RNase H enzymes. The cDNA fragments were then subjected to a series of enzymatic steps that repair the ends, tails the 3’ end with a single ‘A’ base, followed by ligation of the adapters (Refer, Table 1.0 Sample-Index Details). The adapter-ligated products were then purified and enriched using the following thermal conditions: initial denaturation 98°C for 30sec; 14 cycles of - 98°C for 10 sec, 65°C for 75 sec; final extension of 65°C for 5mins. PCR products are then purified and checked for fragment size distribution on High Sensitivity D1000 screen tape (Agilent, Cat no 5067-5584). NEBNext Multiplex Oligos for Illumina 96 Unique Dual Index Primer 96 rxn Set3 (NEB, Cat no E6444S), NEBNext Multiplex Oligos for Illumina 96 Unique Dual Index Primer 96 rxn (NEB, Cat no E6440S) were used for multiplexing. Prepared libraries were quantified using Qubit HS Assay (Invitrogen, Cat no Q32854). The obtained libraries were pooled and diluted to final optimal loading concentration. The pooled libraries were then loaded on to Illumina NovaSeq V1.5 instrument to generate 150 bp Paired end reads.

### RNA seq Analysis

FastQC (Andrews S, 2010) was performed on the fastq files to examine the quality of the reads. Illumina adapters were trimmed using Trimmomatic (0.39) (Bolger, Lohse, & Usadel, 2014) using minimum length of 50 bp and FastQC was rerun to check the quality of the reads. Post trimming, HISAT2 (Kim, Paggi, Park, Bennett, & Salzberg, 2019) was used to align the reads to the mouse genome (version MM10). Alignment for control samples were between 93 to 95% and for mutants they were 96-98%. Post alignment, the Sequence Alignment Files (sam) files were converted to bam files using samtools (Danecek et al., 2021) and using Featurecounts (Liao, Smyth, & Shi, 2014) the counts matrix file was generated. Following the generation of the counts matrix file, DESeq2 (Love, Huber, & Anders, 2014) was run to obtain the normalized gene counts and the differentially expressed genes with the p adjusted value <0.05 and log2 fold change = 0.41 For generation of the heatmap, the ComplexHeatmap package (Gu, Eils, & Schlesner, 2016) from Bioconductor was used. Normalized counts were used to plot a heatmap with the base mean greater than 2000 and log2FC >0.44, p value <0.05.

Raw Fastq files are deposited in the NCBI database with Bioproject accession number:

PRJNA1149579.

### Analysis of scRNAseq data

Single cell data (Di Bella et al., 2021; Zhou et al., 2023) was used to examine the astrocyte transcriptome at E15.5, E18.5, P4 ( (Di Bella et al., 2021; Zhou et al., 2023) and P7 (Zhou et al., 2023), Biorxiv). We used Seurat (version 4.4.0) (Hao et al., 2021) to generate the count matrix file and for further analysis. A scatter plot for nCount_RNA vs nFeature_RNA was used to determine single cells based on RNA content. We performed filtering which included retaining cells with mitochondrial percentage <5 and <5000 genes per cell. PCA was performed on the scaled data and cells were clustered using the FindNeighbors (first 50 principal components) and FindClusters function. UMAP was used to visualize the variation in the cells. For assessing astrocytes, Slc1a3, Aldoc, Aldh1L1 and Gfap were used as shown in feature plots (Supplementary Figure S1B). Astrocytes clusters were subsetted and FindMarkers function was used to examine the differentially expressed genes amongst the clusters with a log2 Fold change threshold of 0.25.

### Pseudo bulk analysis

For analysis shown in Fig, we used the Zhou et al 2023 dataset. Every cell in the control and Olig2 KO was randomly assigned an integer between one and three, and divided into three bins accordingly. The transcriptomic information for cells in each bin was aggregated to create three pseudo-bulk-seq-replicates for control and Olig2 KO conditions. Seurat’s FindMarkers function was used with the test.use argument set to DESeq2, to assess the differential expression of Lhx2 between the two conditions.

### Data analysis

Graphs were generated using GraphPad Prism (Version 9 and 10).

## Notes

### Competing Interest Statement

The authors have declared no competing interest.

## Bibliography

Allen, D. E., Donohue, K. C., Cadwell, C. R., Shin, D., Keefe, M. G., Sohal, V. S., & Nowakowski, T. J. (2022). Fate mapping of neural stem cell niches reveals distinct origins of human cortical astrocytes. Science, 376(6600), 1441–1446.

Allen, N. J., & Eroglu, C. (2017). Cell Biology of Astrocyte-Synapse Interactions. Neuron, 96(3), 697–708.

Ashhad, S., & Narayanan, R. (2016). Active dendrites regulate the impact of gliotransmission on rat hippocampal pyramidal neurons. Proceedings of the National Academy of Sciences of the United States of America, 113(23), E3280–9.

Bardehle, S., Krüger, M., Buggenthin, F., Schwausch, J., Ninkovic, J., Clevers, H., Snippert, H. J., et al. (2013). Live imaging of astrocyte responses to acute injury reveals selective juxtavascular proliferation. Nature Neuroscience, 16(5), 580–586.

Barres, B. A. (2008). The mystery and magic of glia: a perspective on their roles in health and disease. Neuron, 60(3), 430–440.

Bayraktar, O. A., Bartels, T., Holmqvist, S., Kleshchevnikov, V., Martirosyan, A., Polioudakis, D., Ben Haim, L., et al. (2020). Astrocyte layers in the mammalian cerebral cortex revealed by a single-cell in situ transcriptomic map. Nature Neuroscience, 23(4), 500– 509.

Bolger, A. M., Lohse, M., & Usadel, B. (2014). Trimmomatic: A flexible trimmer for Illumina sequence data. Bioinformatics, 30(15), 2114–2120.

Brenner, M., Kisseberth, W. C., Su, Y., Besnard, F., & Messing, A. (1994). GFAP promoter directs astrocyte-specific expression in transgenic mice. The Journal of Neuroscience, 14(3 Pt 1), 1030–1037.

Buffo, A., Rite, I., Tripathi, P., Lepier, A., Colak, D., Horn, A.-P., Mori, T., et al. (2008). Origin and progeny of reactive gliosis: A source of multipotent cells in the injured brain. Proceedings of the National Academy of Sciences of the United States of America, 105(9), 3581–3586.

Chen, Y., Miles, D. K., Hoang, T., Shi, J., Hurlock, E., Kernie, S. G., & Lu, Q. R. (2008). The basic helix-loop-helix transcription factor olig2 is critical for reactive astrocyte proliferation after cortical injury. The Journal of Neuroscience, 28(43), 10983–10989.

Clavreul, S., Abdeladim, L., Hernández-Garzón, E., Niculescu, D., Durand, J., Ieng, S.-H., Barry, R., et al. (2019). Cortical astrocytes develop in a plastic manner at both clonal and cellular levels. Nature Communications, 10(1), 4884.

Danecek, P., Bonfield, J. K., Liddle, J., Marshall, J., Ohan, V., Pollard, M. O., Whitwham, A., et al. (2021). Twelve years of SAMtools and BCFtools. GigaScience, 10(2).

Di Bella, D. J., Habibi, E., Stickels, R. R., Scalia, G., Brown, J., Yadollahpour, P., Yang, S. M., et al. (2021). Molecular logic of cellular diversification in the mouse cerebral cortex. Nature, 595(7868), 554–559.

Escartin, C., Galea, E., Lakatos, A., O’Callaghan, J. P., Petzold, G. C., Serrano-Pozo, A., Steinhäuser, C., et al. (2021). Reactive astrocyte nomenclature, definitions, and future directions. Nature Neuroscience, 24(3), 312–325.

Ganat, Y. M., Silbereis, J., Cave, C., Ngu, H., Anderson, G. M., Ohkubo, Y., Ment, L. R., et al. (2006). Early postnatal astroglial cells produce multilineage precursors and neural stem cells in vivo. The Journal of Neuroscience, 26(33), 8609–8621.

Ge, W.-P., Miyawaki, A., Gage, F. H., Jan, Y. N., & Jan, L. Y. (2012). Local generation of glia is a major astrocyte source in postnatal cortex. Nature, 484(7394), 376–380.

Gu, Z., Eils, R., & Schlesner, M. (2016). Complex heatmaps reveal patterns and correlations in multidimensional genomic data. Bioinformatics, 32(18), 2847–2849.

Hao, Y., Hao, S., Andersen-Nissen, E., Mauck, W. M., Zheng, S., Butler, A., Lee, M. J., et al. (2021). Integrated analysis of multimodal single-cell data. Cell, 184(13), 3573–3587.

Hasel, P., Rose, I. V. L., Sadick, J. S., Kim, R. D., & Liddelow, S. A. (2021). Neuroinflammatory astrocyte subtypes in the mouse brain. Nature Neuroscience, 24(10), 1475–1487.

John Lin, C.-C., Yu, K., Hatcher, A., Huang, T.-W., Lee, H. K., Carlson, J., Weston, M. C., et al. (2017). Identification of diverse astrocyte populations and their malignant analogs. Nature Neuroscience, 20(3), 396–405.

Kim, D., Paggi, J. M., Park, C., Bennett, C., & Salzberg, S. L. (2019). Graph-based genome alignment and genotyping with HISAT2 and HISAT-genotype. Nature Biotechnology, 37(8), 907–915.

Lanjakornsiripan, D., Pior, B.-J., Kawaguchi, D., Furutachi, S., Tahara, T., Katsuyama, Y., Suzuki, Y., et al. (2018). Layer-specific morphological and molecular differences in neocortical astrocytes and their dependence on neuronal layers. Nature Communications, 9(1), 1623.

Liao, Y., Smyth, G. K., & Shi, W. (2014). featureCounts: an efficient general purpose program for assigning sequence reads to genomic features. Bioinformatics, 30(7), 923– 930.

Love, M. I., Huber, W., & Anders, S. (2014). Moderated estimation of fold change and dispersion for RNA-seq data with DESeq2. Genome Biology, 15(12), 550.

MacVicar, B. A., & Newman, E. A. (2015). Astrocyte regulation of blood flow in the brain. Cold Spring Harbor Perspectives in Biology, 7(5), a020388.

Madisen, L., Zwingman, T. A., Sunkin, S. M., Oh, S. W., Zariwala, H. A., Gu, H., Ng, L. L., et al. (2010). A robust and high-throughput Cre reporting and characterization system for the whole mouse brain. Nature Neuroscience, 13(1), 133–140.

Mangale, V. S., Hirokawa, K. E., Satyaki, P. R. V., Gokulchandran, N., Chikbire, S., Subramanian, L., Shetty, A. S., et al. (2008). Lhx2 selector activity specifies cortical identity and suppresses hippocampal organizer fate. Science, 319(5861), 304–309.

Ma, S., Kwon, H. J., & Huang, Z. (2012). A functional requirement for astroglia in promoting blood vessel development in the early postnatal brain. Plos One, 7(10), e48001.

de Melo, J., Miki, K., Rattner, A., Smallwood, P., Zibetti, C., Hirokawa, K., Monuki, E. S., et al. (2012). Injury-independent induction of reactive gliosis in retina by loss of function of the LIM homeodomain transcription factor Lhx2. Proceedings of the National Academy of Sciences of the United States of America, 109(12), 4657–4662.

Miller, F. D., & Gauthier, A. S. (2007). Timing is everything: making neurons versus glia in the developing cortex. Neuron, 54(3), 357–369.

Mishra, P., & Narayanan, R. (2020). Heterogeneities in intrinsic excitability and frequency-dependent response properties of granule cells across the blades of the rat dentate gyrus. Journal of Neurophysiology, 123(2), 755–772.

Mishra, P., & Narayanan, R. (2022). Conjunctive changes in multiple ion channels mediate activity-dependent intrinsic plasticity in hippocampal granule cells. iScience, 25(3), 103922.

Morel, L., Chiang, M. S. R., Higashimori, H., Shoneye, T., Iyer, L. K., Yelick, J., Tai, A., et al. (2017). Molecular and functional properties of regional astrocytes in the adult brain. The Journal of Neuroscience, 37(36), 8706–8717.

Narayanan, R., & Johnston, D. (2007). Long-term potentiation in rat hippocampal neurons is accompanied by spatially widespread changes in intrinsic oscillatory dynamics and excitability. Neuron, 56(6), 1061–1075.

Narayanan, R., & Johnston, D. (2008). The h channel mediates location dependence and plasticity of intrinsic phase response in rat hippocampal neurons. The Journal of Neuroscience, 28(22), 5846–5860.

Otsu, Y., Couchman, K., Lyons, D. G., Collot, M., Agarwal, A., Mallet, J.-M., Pfrieger, F. W., et al. (2015). Calcium dynamics in astrocyte processes during neurovascular coupling. Nature Neuroscience, 18(2), 210–218.

Sofroniew, M. V., & Vinters, H. V. (2010). Astrocytes: biology and pathology. Acta Neuropathologica, 119(1), 7–35.

Sofroniew, M. V. (2005). Reactive astrocytes in neural repair and protection. The Neuroscientist, 11(5), 400–407.

Somjen, G. G. (1988). Nervenkitt: notes on the history of the concept of neuroglia. Glia, 1(1), 2–9.

Stogsdill, J. A., Ramirez, J., Liu, D., Kim, Y. H., Baldwin, K. T., Enustun, E., Ejikeme, T., et al. (2017). Astrocytic neuroligins control astrocyte morphogenesis and synaptogenesis. Nature, 551(7679), 192–197.

Subramanian, L., Sarkar, A., Shetty, A. S., Muralidharan, B., Padmanabhan, H., Piper, M., Monuki, E. S., et al. (2011). Transcription factor Lhx2 is necessary and sufficient to suppress astrogliogenesis and promote neurogenesis in the developing hippocampus. Proceedings of the National Academy of Sciences of the United States of America, 108(27), E265–74.

Tabata, H., Sasaki, M., Agetsuma, M., Sano, H., Hirota, Y., Miyajima, M., Hayashi, K., et al. (2022). Erratic and blood vessel-guided migration of astrocyte progenitors in the cerebral cortex. Nature Communications, 13(1), 6571.

Tani, H., Dulla, C. G., Farzampour, Z., Taylor-Weiner, A., Huguenard, J. R., & Reimer, R. J. (2014). A local glutamate-glutamine cycle sustains synaptic excitatory transmitter release. Neuron, 81(4), 888–900.

Weng, Q., Wang, J., Wang, J., He, D., Cheng, Z., Zhang, F., Verma, R., et al. (2019). Single-Cell Transcriptomics Uncovers Glial Progenitor Diversity and Cell Fate Determinants during Development and Gliomagenesis. Cell Stem Cell, 24(5), 707-723.e8.

Zhang, Y., Chen, K., Sloan, S. A., Bennett, M. L., Scholze, A. R., O’Keeffe, S., Phatnani, H. P., et al. (2014). An RNA-sequencing transcriptome and splicing database of glia, neurons, and vascular cells of the cerebral cortex. The Journal of Neuroscience, 34(36), 11929–11947.

Zhou, J., Vitali, I., Roig Puiggros, S., Javed, A., Jabaudon, D., Mayer, C., & Bocchi, R. (2023). Dual lineage origins of neocortical astrocytes. BioRxiv.

