## Supplementary Figures and Information for "Transcription factor LHX2 suppresses astrocyte proliferation in the postnatal mammalian cerebral cortex"

#### SUPPLEMENTARY FIGURE S1

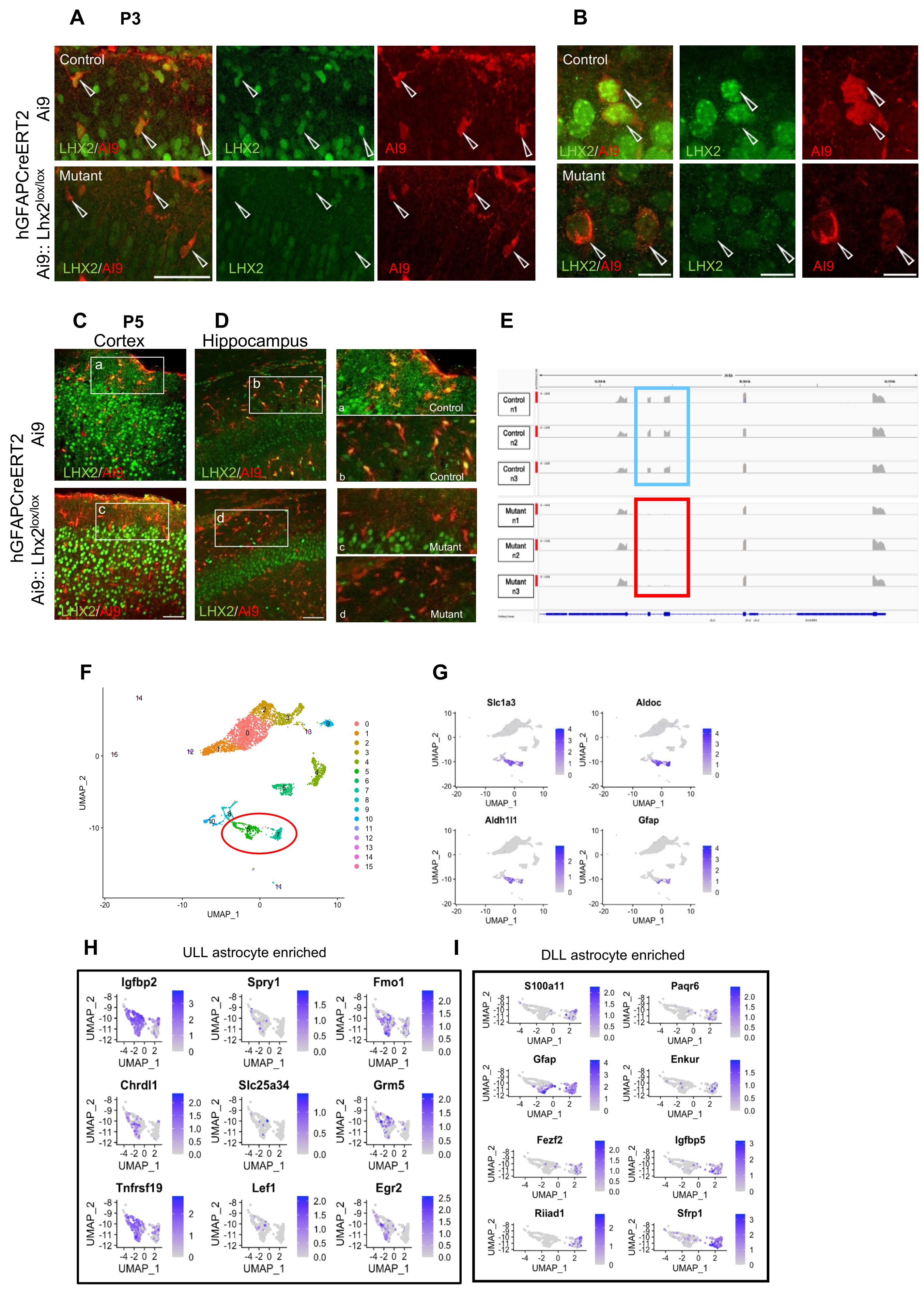

**SUPPLEMENTARY FIGURE S2**

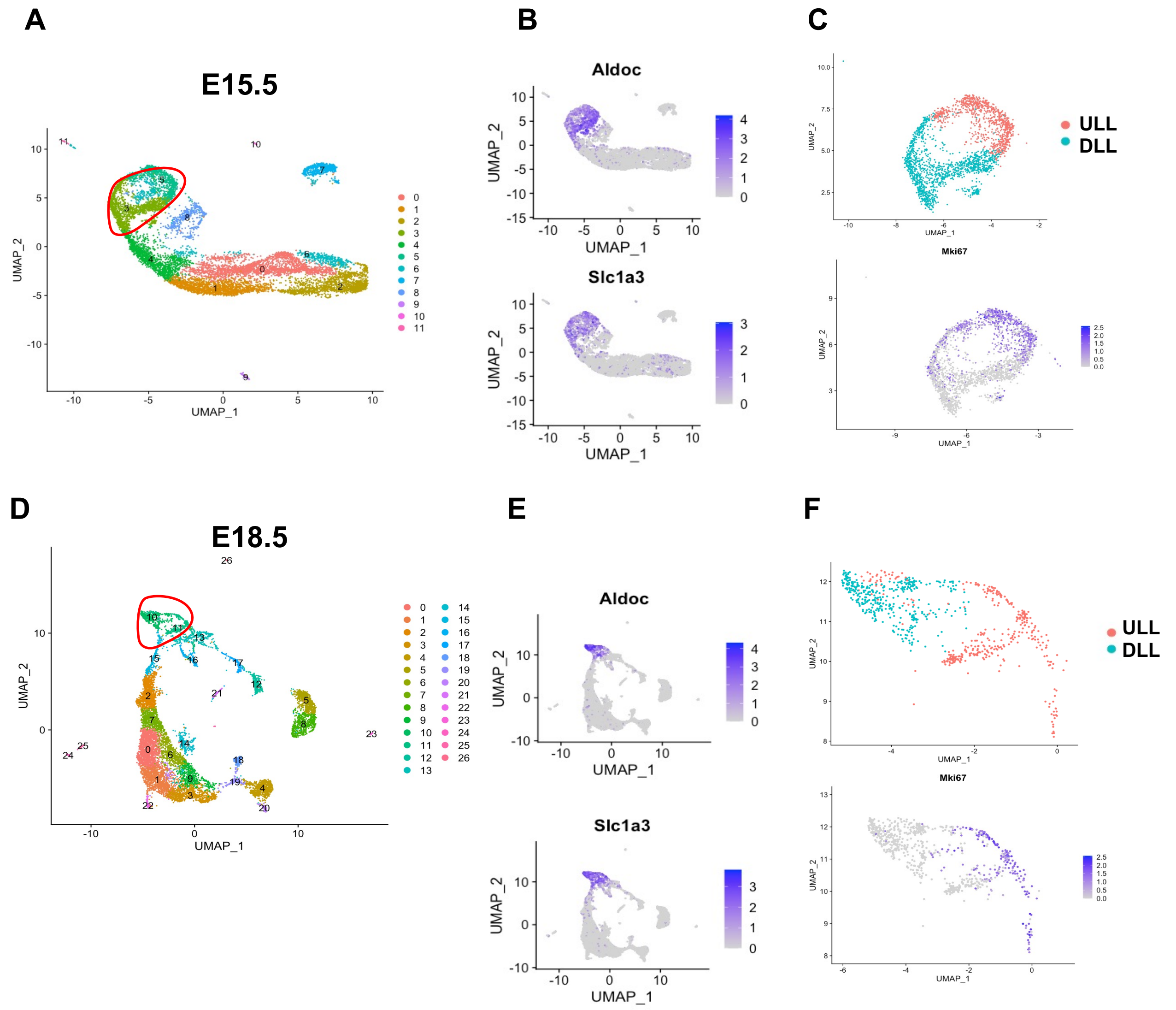

### SUPPLEMENTARY FIGURE S3

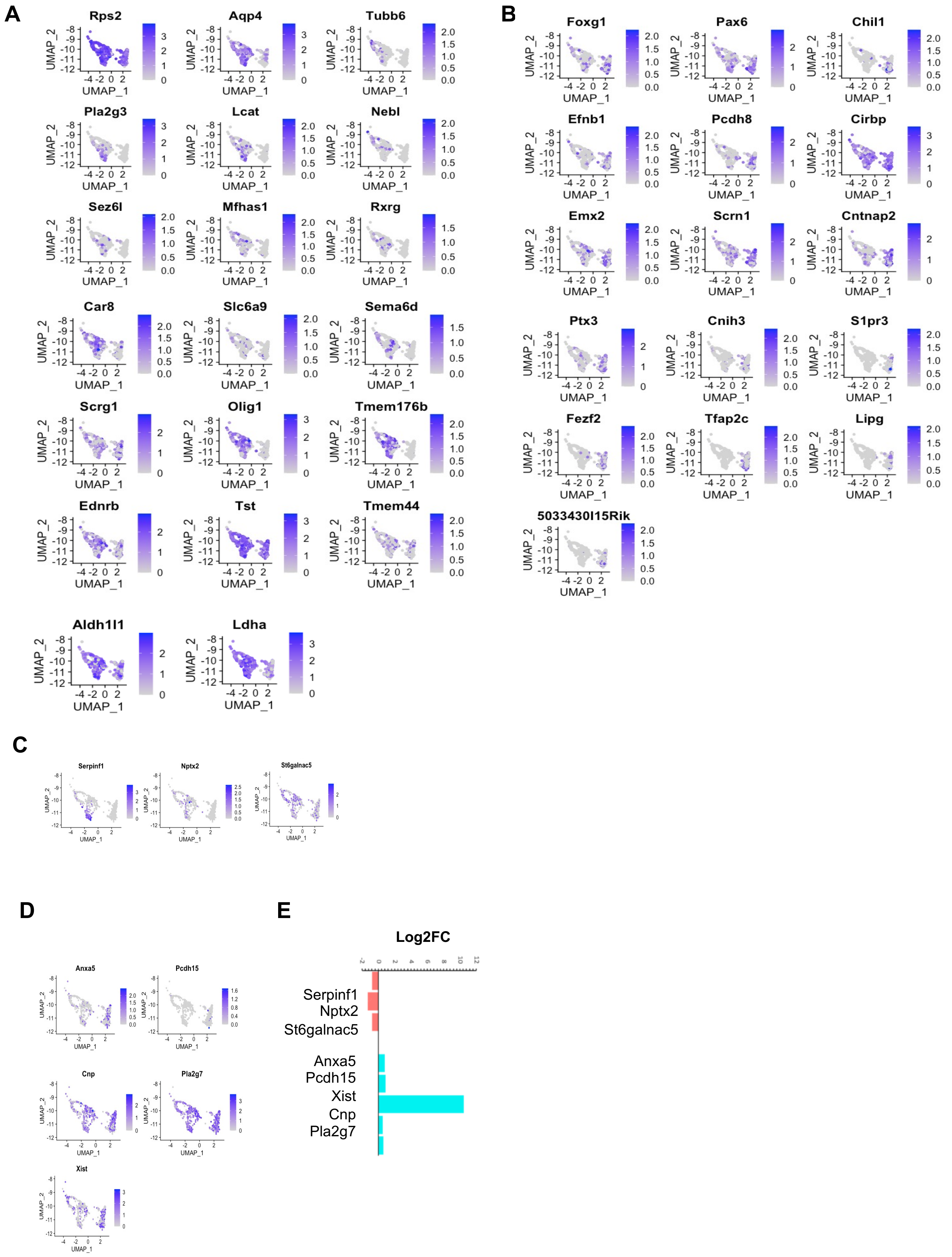

### SUPPLEMENTARY FIGURE S4

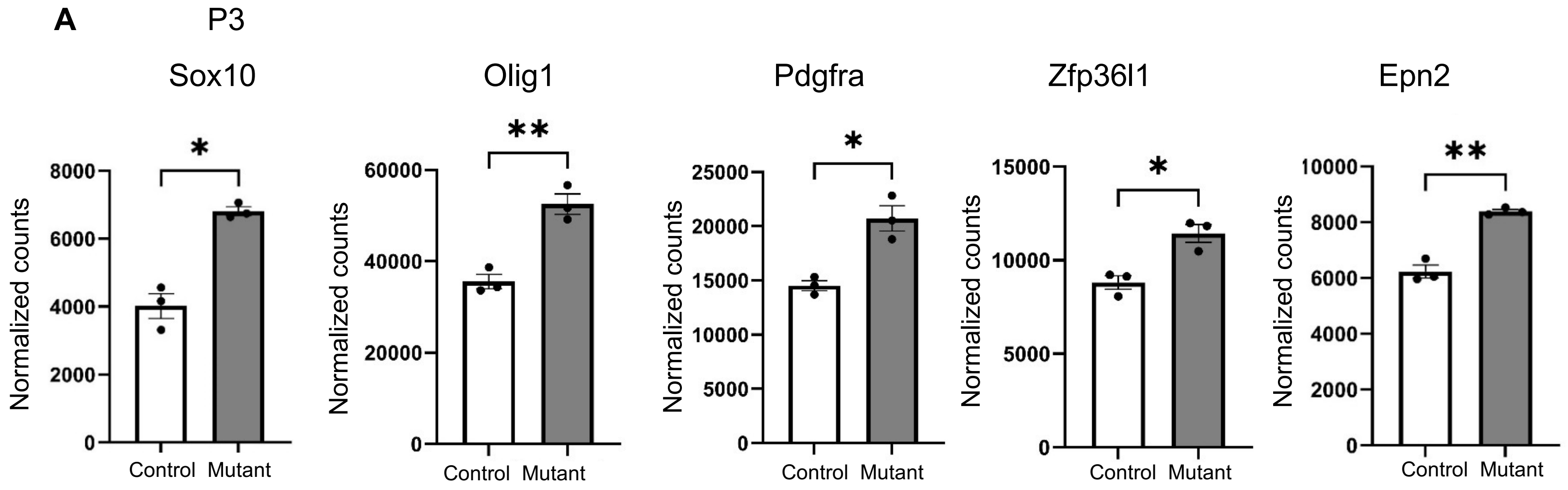


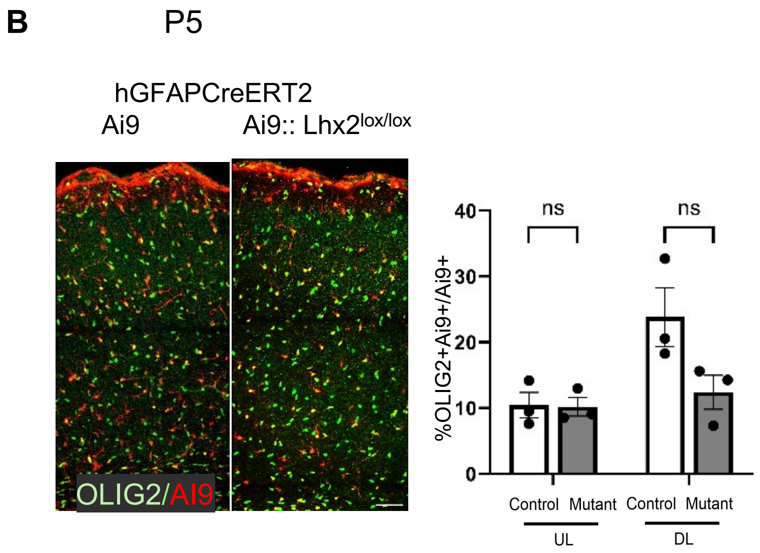

### SUPPLEMENTARY FIGURE S5

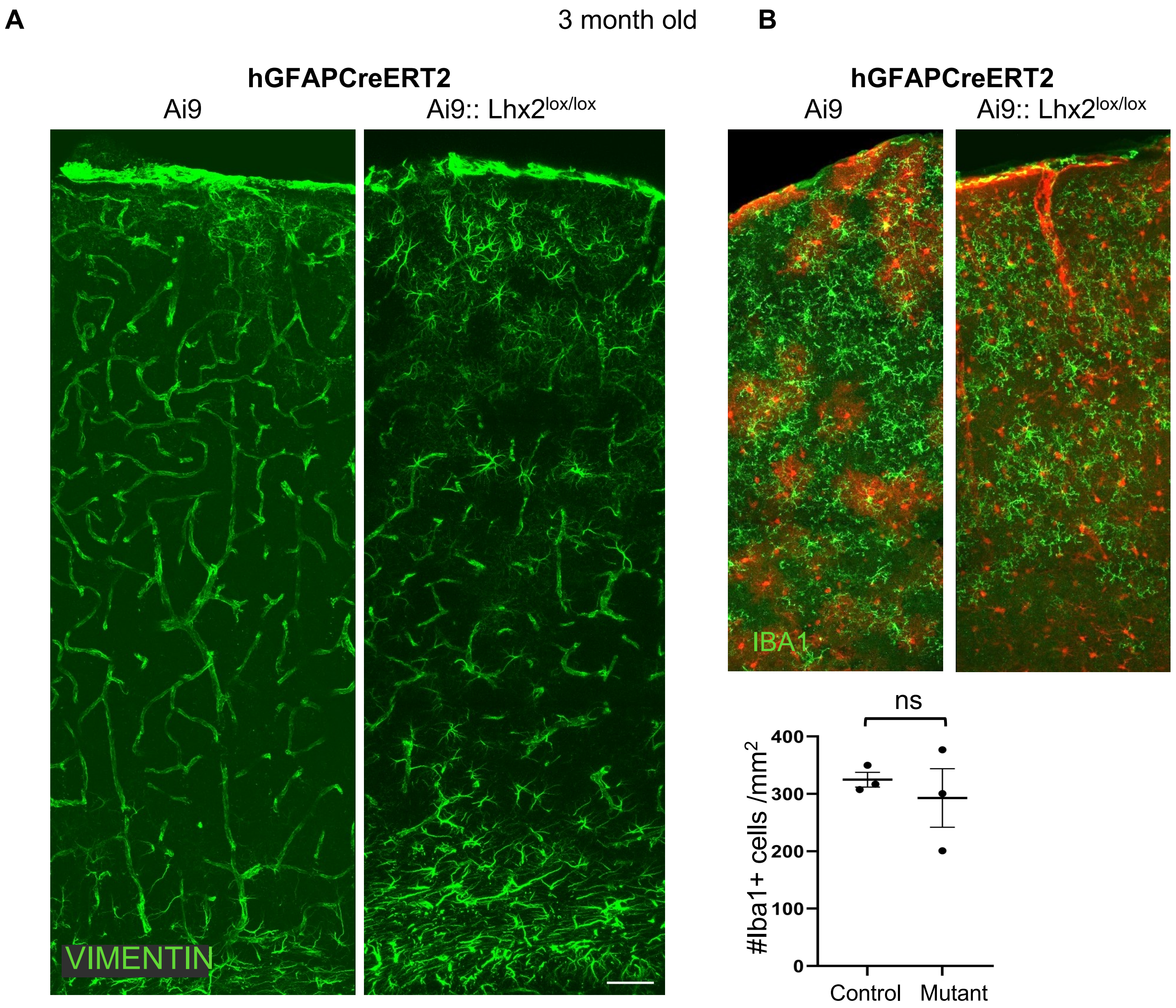

**SUPPLEMENTARY FIGURE S6**

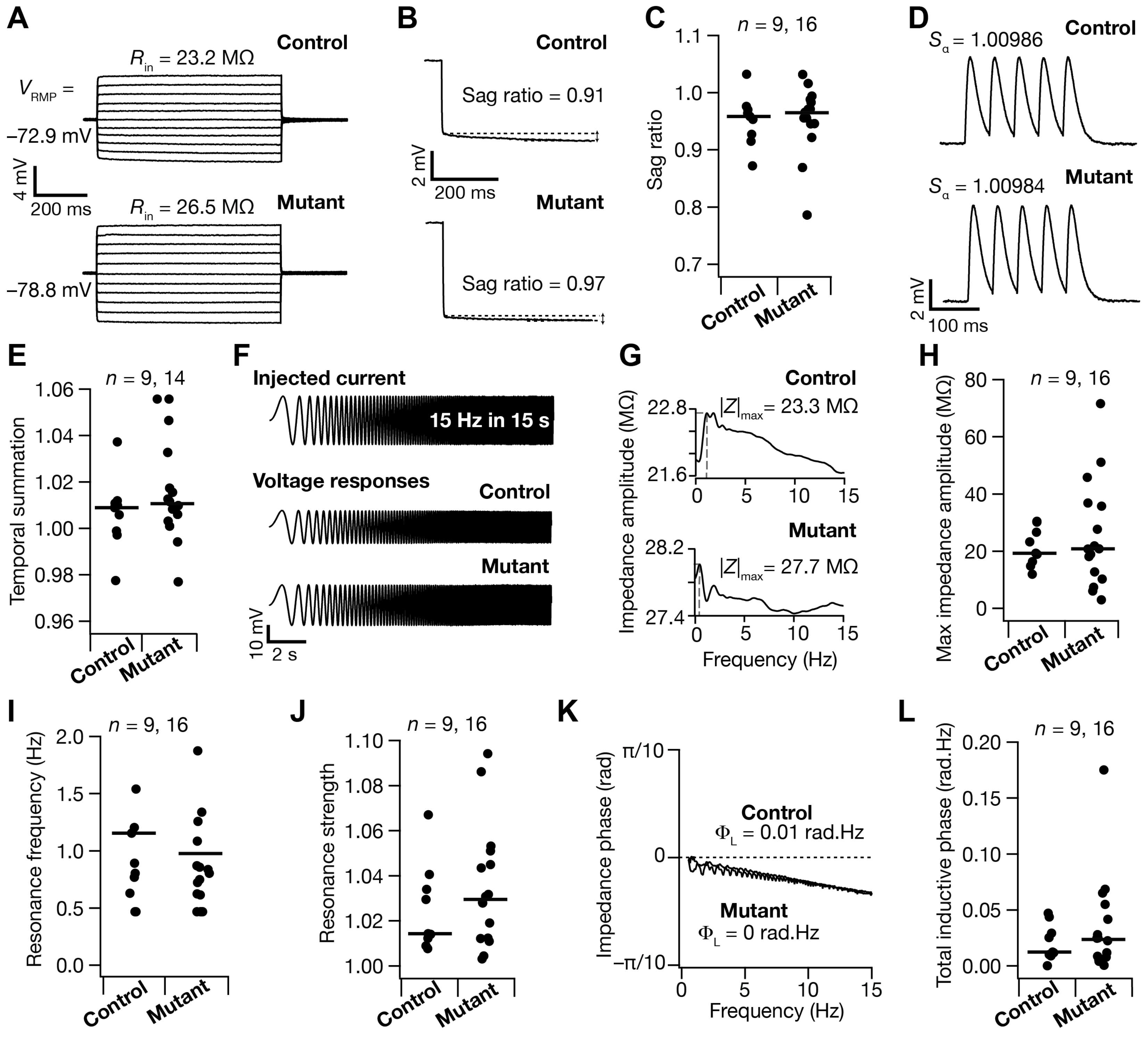

#### APPENDIX

| Antibody and drugs | Company, catalog number and dilution used | Lot number |
|---|---|---|
| Mouse Sox2 | Invitrogen MA1-014, 1:100 | VB294349 |
| Rabbit Lhx2 | Merck Abe1402, 1:200 | 3030529 |
| Mouse RFP | Invitrogen MA515257, 1:200 | 1U00903 |
| Rabbit SOX9 | Invitrogen PA5-81966, 1:200 | YK4128713 |
| Rabbit KI67 | Invitrogen MA5-14520, 1:200 | XH3680491 |
| Mouse ALDH1L1 | Invitrogen 13-9595-82, 1:50 | 2549557 |
| Rabbit VIMENTIN | Abcam ab92547, 1:750 | Epr3776 |
| Rabbit GFAP | Sigma G9269, 1:200 | O25M4843V |

**Supplementary Figure S1**

(A -D) Immunostaining to examine the presence of LHX2 protein after tamoxifen administration at P1 to hGFAPCreERT2:: Ai9 (control) and hGFAPCreERT2:: Ai9::Lhx2$^{lox/lox}$(mutant) animals.

(A, B) At P3, Ai9+ cells (open arrowheads) display LHX2 immunoreactivity in control but weak to undetectable signal in mutant brains.

(C, D) At P5, both in the neocortex (C) and in the hippocampus (D), LHX2 immunoreactivity is seen in Ai9+ cells (open arrowheads) in control brains but is undetectable in mutants. Regions in white boxes marked (a-d) are presented at high magnification in adjacent panels.

(E) IGV browser view of the *Lhx2* genomic region showing mapped reads corresponding to control and mutant RNAseq data from FACS sorted Ai9+ cells at P3 showing the presence of exon 2 and 3 in control (blue box) and absence in the mutant samples (red box).

(F) UMAP of P4 scRNA seq data from Di Bella et al 2021 (G) Markers used to subset astrocytes

(H) Feature plots of known UL and (I) DL enriched astrocyte markers

Scale bars in A, C and D are 50µm; Scale bars in B:10µm

**Supplementary Figure S2**

A and D) UMAP of E15.5 and E18.5 scRNA seq data from Di Bella et al 2021

B and E) Markers used to subset astrocytes

C and F) Two clusters within astrocyte population representing ULL and DLL signatures

**Supplementary Figure S3**

(A) (B) and (C) Feature plots showing expression of dysregulated genes upon Lhx2 loss enriched in ULL astrocytes and in (D) and (E) DLL astrocytes (F) Log2 Fold change of 3 of the genes enriched in ULL cluster (Salmon) and were downregulated upon Lhx2 loss and the 5 genes enriched in the DLL cluster (Cyan) and that were upregulated.

**Supplementary Figure S4**

(A) Normalised counts of OPC markers *Sox10, Olig1, Pdgfra, Zfp36l1* and *Epn2* were examined from the P3 RNA seq data obtained from hGFAPCreERT2:: Ai9 (control) and hGFAPCreERT2:: Ai9::Lhx2$^{lox/lox}$(mutant) Ai9+ cells. The expression of each gene is increased in the mutant. Statistical test: Unpaired t test with Welch's correction. * $p<0.05$, ** $p<0.005$. (B) OLIG2 immunostaining and quantification at P5 shows no difference in the OLIG2+ AI9 + fraction of Ai9+ cells between the two groups in both UL (p=0.99) and DL (p=0.12). Statistical test: Two-way ANOVA with Sidak's multiple comparison test; Control: n= 1998 Ai9 cells; Mutant: n= 1683 cells; N= 3 independent biological replicates. Error bars depict SEM. ns: not significant

**Supplementary Figure S5**

(A, B) VIMENTIN and (IBA1) immunostaining in 3-month-old control and mutant brains. The VIMENTIN image (A) is a single-channel presentation of data that is quantified in Figure 4A, B, for ease of visualization of the increased VIMENTIN+ astrocytes in the grey matter. In contrast, IBA1 staining (B) reveals no difference in distribution between control and mutant brains. Quantification of IBA1+ cells / $mm^2$ (Control: 324 cells± 13, Mutant 275 ± 52, p = 0.4, Unpaired t test) (C) *Sox10* expression in Ai9+ cells is increased upon loss of *Lhx2* (p=0.01, Unpaired t test with Welch's correction). Data from normalised counts obtained from RNA seq (Figure 2D)

**Supplementary Figure S6**

Electrophysiological characterization of control and mutant astrocytes. (A) Example voltage responses of control (top) and mutant (bottom) astrocytes to pulse current injections ranging from –250 pA to +250 pA in steps of 50 pA, used for computing input resistance ($R_{in}$). (B) Traces showing voltage response to a 250-pA hyperpolarizing pulse current for the example control (top) and mutant (bottom) astrocytes. Sag ratio was computed as the ratio between the initial peak and the steady-state voltage deflection. (C) Beeswarm plot of Sag ratio recorded from all control and mutant astrocytes. $p$=0.6 for Sag. (D) Train of five excitatory postsynaptic potentials (α-EPSPs) recorded from example control (top) and mutant (bottom) astrocytes. Temporal summation ($S_\alpha$) was calculated as the ratio between the last and the first EPSP amplitudes. (E) Beeswarm plot of $S_\alpha$ recorded from all control and mutant astrocytes. $p$=0.89 for $S_\alpha$. (F) *Top*, Chirp current stimulus used for characterizing frequency-dependent characteristics of cortical astrocytes. *Bottom*, Voltage responses of the example control and mutant astrocytes to the chirp current stimulus. (G) Plots showing impedance amplitude profiles of control and mutant astrocytes with resonance frequency ($f_R$=1.15 Hz for control and 0.87 Hz for mutant examples) defined as the frequency at which maximum impedance ($|Z|_{max}$) was achieved. (H–J) Beeswarm plots of maximum impedance amplitude ($|Z|_{max}$), resonance frequency ($f_R$), and resonance strength ($Q$) calculated for all control and mutant astrocytes. $p$=0.93 for $|Z|_{max}$, $p$=0.6 for $f_R$ and $p$=0.73 for $Q$. (K) Plot showing impedance phase profile control and mutant astrocytes. Note that there was no pronounced shift in impedance phase from zero, over the entire range of frequencies measured. (L) Beeswarm plot of total inductive phase ($\Phi_L$) calculated for all control and mutant astrocytes respectively. $p$=0.98 for $\Phi_L$. All illustrative example traces in the figure were taken from a single control and a single mutant astrocyte. The thin lines in all beeswarm plots indicate the respective median values. All $p$ values correspond to the Wilcoxon rank sum test.

**Supplementary Movie 1 and 2**: Videos showing presence of VIMENTIN in control and mutant brain slices.

**Supplementary Table 1:**

Differentially expressed genes between hGFAPCreERT2:: Ai9 (control) and hGFAPCreERT2:: Ai9::Lhx2$^{lox/lox}$(mutant) conditions obtained from P3 RNA sequencing of Ai9+ sorted cells.

**Supplementary Table 2:**

Differentially expressed genes between Upper Layer Like (ULL) cluster and Deep Layer like (DLL) analysed from Di Bella et al 2021 dataset (avg log 2FC (0.25)

#### Appendix

| **Antibodies used** | **Company, catalog number and dilution used** | **Lot number** |
|---|---|---|
| Mouse Anti Sox2 | Invitrogen MA1-014, 1:100 | VB294349 |
| Rabbit Anti Lhx2 | Merck Abe1402, 1:200 | 3030529 |
| Mouse Anti RFP | Invitrogen MA515257, 1:200 | 1U00903 |
| Rabbit Anti SOX9 | Invitrogen PA5-81966, 1:200 | YK4128713 |
| Rabbit Anti KI67 | Invitrogen MA5-14520, 1:200 | XH3680491 |
| Mouse Anti ALDH1L1 | Invitrogen 13-9595-82, 1:50 | 2549557 |
| Rabbit Anti VIMENTIN | Abcam ab92547, 1:750 | Epr3776 |

| | | |
|---|---|---|
| Rabbit Anti GFAP | Sigma G9269, 1:200 | O25M4843V |
| Goat Anti IBA1 | Abcam [ab5076], 1:1000 | GR254159-6 |
